# Loss of ANT1 Increases Fibrosis and Epithelial Cell Senescence in Idiopathic Pulmonary Fibrosis

**DOI:** 10.1101/2022.09.09.507271

**Authors:** Jennifer C. Boatz, Justin Sui, Qianjiang Hu, Xiaoyun Li, Yingze Zhang, Melanie Königshoff, Corrine R. Kliment

**Author notes:** Corresponding Author: Corrine R. Kliment, W1254, Biomedical Science Tower, 200 Lothrop Street, Pittsburgh, PA 15213, Office. Co-first authors (Jennifer C. Boatz, Justin Sui). **Author contribution’s:** Conceived the experiments: JCB and CRK; Collection of samples: JBC, JS, and CRK; Performed the experiments: CRK, JCB, QH, XL; Data analysis: CRK, JCB, QH; Wrote the paper and figure creation: JS, JCB and CRK; Critical review and approval: YZ, MK, and CRK. All authors reviewed the manuscript and approved the final version prior to submission.

## Abstract

Idiopathic Pulmonary Fibrosis (IPF) is an interstitial lung disease characterized by progressive lung scarring and remodeling. Although treatments exist that slow disease progression, IPF is irreversible and there is no cure. Cellular senescence, a major hallmark of aging, has been implicated in IPF pathogenesis, and mitochondrial dysfunction is increasingly recognized as a driver of senescence. Adenine nucleotide translocases (ANTs) are abundant mitochondrial ATP-ADP transporters critical for regulating cell fate and maintaining mitochondrial function. We sought to determine how alterations in ANTs influence cellular senescence in pulmonary fibrosis. We found *SLC25A4* (ANT1) and *SLC25A5* (ANT2) expression is reduced in the lungs of IPF patients and particularly within alveolar type II cells by single cell RNA sequencing. Loss of ANT1 by siRNA in lung epithelial cell lines resulted in increased senescence markers such as beta-galactosidase staining and p21 by Western Blot and RT-qPCR. Bleomycin treated ANT1 knockdown cells also had increased senescence markers when compared to bleomycin treated control cells. Global loss of ANT1 resulted in worse lung fibrosis and increased senescence in the bleomycin and asbestos-induced mouse models of pulmonary fibrosis. This data supports the concept that loss of ANT1 drives IPF pathogenesis through mitochondrial dysfunction associated cellular senescence (MiDaS). In summary, loss of ANT1 induces cellular senescence, leading to abnormal tissue remodeling and enhanced lung fibrosis in IPF. Modulation of ANTs presents a new therapeutic avenue that may alter cellular senescence pathways and limit pulmonary fibrosis.

## Introduction

Idiopathic pulmonary fibrosis (IPF) is a deadly and progressive lung disease characterized by irreversible scarring, or fibrosis, of the lungs, remodeling and dysregulated epithelial repair (1). Current treatments, including anti-fibrotic agents have limited efficacy on reversing or preventing disease (2). Studies have demonstrated that, in addition to a loss of alveolar epithelial cells in the IPF (3), there are complex changes that occur in the remaining epithelial cells in the lung. These alveolar epithelial cells undergo functional reprogramming and acquire “transitional” cell states not typically seen in the healthy lung (3,4). A deeper understanding of the mechanisms that result in epithelial cell loss and reprogramming is needed to develop targeted therapeutics that can stop or even reverse disease progression.

One important component of IPF pathogenesis is cellular senescence (5), a permanent state of cell cycle arrest that plays a role in several aging related diseases (6,7). Senescent cells are metabolically active cells that often resist apoptosis and accumulate in organs as part of an aging-associated stress response to stimuli. Mitochondrial dysfunction has been increasingly recognized as an important inducer of senescence (8), however, additional research is necessary to define this process. Changes in mitochondrial structure and function have been implicated in aging and IPF, and have primarily been observed in lung epithelial cells, fibroblasts, and macrophages (9–11). Mitochondrial dysfunction associated senescence (MiDAS) has recently been described in IPF (8,12,13). The cellular factors driving MiDAS and fibrosis progression remain unclear and warrant more investigation.

The gene expression profile of senescent cells is characteristically altered compared to non-senescent cells (14), most notably with upregulation of cell cycle inhibitors such as p16^INK4a^ and p21^CIP1/WAF1^, tumor suppressor protein p53 and increased lysosomal beta-galactosidase (β-gal) activity. Senescent cells also secrete a milieu of signaling molecules, including proinflammatory chemokines and growth factors, collectively known as the senescence-associated secretory phenotype (SASP) (15) that may propagate inflammation and tissue remodeling. Upregulation of these senescence markers and SASP have been identified in epithelial cells and fibroblasts in lung fibrosis (5,16). The clearance of these senescent cells and SASP inhibition have both been found to enhance pulmonary function in mouse models (17,18). Thus, the identification of pharmacological treatments that target senescent cells, MiDAS or SASP suppression might be promising avenues of IPF treatment.

Adenine nucleotide translocases (human paralogs ANT1-4) are mitochondrial ADP/ATP transporters that are critical for mitochondrial bioenergetics, reactive oxygen species (ROS) production, and regulation of cell fate (19,20). ANT1 and ANT2 are expressed in the human lung (21). Several mutations in ANTs, particular in ANT1, have been described in human mitochondrial disease (22). Null and point mutations for ANT1 result in mitochondrial myopathy, hypertrophic cardiomyopathy, lactic acidosis, and ophthalmoplegia (22). Despite evidence that ANT1 is important for maintaining mitochondrial function, the role of ANT1 in cellular senescence, MiDAS and lung disease has not yet been described. By leveraging human lung tissue and the bleomycin and asbestos injury models in mice, we sought to determine how alterations in ANT1 influence cellular senescence in pulmonary fibrosis.

## Materials and methods

See **Supplemental Data** for expanded method.

### Human lung tissue studies

Human lung tissue samples were obtained from the Tissue Core at the University of Pittsburgh (supported by P30 DK072506, NIDDK and the CFF RDP). Control lung samples originated from lungs deemed unsuitable for organ transplantation. All IPF samples were from lungs explanted from IPF patients that had undergone lung transplantation under an approved protocol (STUDY18100070). Genomic data was obtained from the Lung Genomics Research Consortium (LGRC) (GEO GSE47460; http://www.lung-genomics.org/) (23) using tissue samples and clinical data collected through the Lung Tissue Research Consortium (LTRC; http://www.ltrcpublic.com/). Single cell RNA sequencing data is publicly available at NCBI’s Gene Expression Omnibus (GSE190889) and a GitHub repository (https://github.com/KonigshoffLab/GPR87_IPF_2022). See the Extended Methods for additional information.

### Bleomycin and asbestos mouse models

All animal studies were approved by the University of Pittsburgh IACUC. ANT1-null mice were a gift from Douglas Wallace (University of Pennsylvania). Male and female mice (10-12 weeks old, n=6-10 per group) were treated with either intratracheal bleomycin sulfate (0.05U per 25g mouse with dose adjustments for weight) or saline control or intratracheal crocidolite asbestos or titanium dioxide (inert control particle) (0.1mg in saline) (24). Mice were euthanized at 28 days post-treatment. Bronchoalveolar lavage (BAL) was collected, and lungs were inflated with 10% formalin at a pressure of 25 cm^2^ H_2_O and fixed in formalin for 24 hr before paraffin embedding or flash frozen in liquid nitrogen for protein analysis, RNA isolation or hydroxyproline quantification. Cytokine concentrations in BALs were measured by Luminex Cytokine Multiplex Panel (R&D Systems) according to the manufacturer’s protocols.

### Immunocytochemistry of mouse and human lung tissue

Immunofluorescent staining of lung tissue was completed as previously described (21). IPF patients and normal control lungs were fixed with 10% formalin and embedded in paraffin. Human and mouse lung sections were stained for: ANT1 (Abcam #ab102032, 1:500), ANT2 (Abcam #192410, 1:100), and secondary Alexa fluorophore antibodies (Molecular Probes). Control sections were stained with non-immune rabbit IgG (#2729P, Cell signaling). Images were acquired on a Nikon A1R confocal microscope.

### Human cell culture and targeted gene suppression

Human cell lines utilized include the bronchial epithelial cell lines HBEC3-KT (gift from John Minna) and BEAS-2B (ATCC) and alveolar epithelial cells A549s (ATCC). Epithelial cell lines were transfected with 150 nM siRNA ON-TARGETplus smart pools (Dharmacon: ANT1 (*SLC25A4*, #L-007485-00-0005), ANT2 (*SLC25A5*, #L-007486-02-0005), non-targeting control pool (#D-001810-10-05) using Lipofectamine 3000 (Invitrogen) according to manufacturer’s protocol. To induce senescence, A549 cells were treated with 10 μg/mL bleomycin over 2 days. RT-qPCR, western blot and beta-galactosidase staining were completed as described in the supplemental methods.

### Statistics

Mean densitometry (ImageLab Software) and all other quantitative data (mean +/- SEM) are normalized to appropriate control groups. Statistical analyses were completed using GraphPad Prism 9.3. Data are assessed for sample distribution. If samples are normally distributed, then data are analyzed using ANOVA with Fisher’s LSD post-test. If the data are not normally distributed, nonparametric analyses, including Kruskal-Wallis, Tukey’s, and/or Mann-Whitney are used.

## Results

### ANT expression is reduced in the lungs of patients with IPF

To determine whether ANTs play a role in IPF pathogenesis, we compared whole lung tissue gene expression for *SLC25A4* (ANT1) and *SLC25A5* (ANT2) in lung tissue from patients with IPF (**Figure 1A**) in an Illumina microarray dataset from the LGRC. We found that gene expression for *SLC25A4* (*P value < 0.0001) and *SLC25A5* (*P value < 0.0001) was significantly reduced in the lungs of patients with IPF (n=255) compared to healthy controls (n=137) (**Figure 1A**). To validate these findings, we confirmed that *SLC25A4* and *SLC25A5* mRNA levels are reduced in lung tissue of patients with IPF from a separate cohort of IPF (n=40) and normal control lung (n=29) by real time PCR (RT-qPCR) (**Figure 1B**, *P value = 0.0029 for *SLC25A4* and *P< 0.0001 for *SLC25A5*). To further localize the expression of ANT proteins in the lung *in situ*, we stained human lung sections from 3 patients with IPF and 3 healthy controls for ANT1 and ANT2 proteins (**Figure 1C, D, Supplemental Figure E1**). Importantly, we found that ANT1 and ANT2 protein expression was moderately reduced in alveolar cells in fibrotic alveolar tissue from the IPF group compared to the alveolar tissue from the controls. Alveolar epithelial cells are a key cell type that are affected in IPF. We next utilized a human single cell RNA sequencing (scRNAseq) dataset of EpCAM+ lung epithelial cells, isolated from healthy control and IPF lungs to obtain a more comprehensive understanding of how *SCL25A4* and *SLC25A5* transcript levels change between different epithelial cell types. Gene expression of *SCL25A4* and *SLC25A5* were markedly reduced in alveolar type I (AT1) and type II (AT2) cells from IPF lungs compared to normal lungs (**Figure 1E**). Together, these observations show that ANT1 and ANT2 expression is reduced in fibrotic lung tissue, particularly in AT1 and AT2 cells. This supports the notion that loss of ANTs may play an important role in alveolar epithelial cells and might contribute to the pathogenesis of IPF.

**Figure 1:**
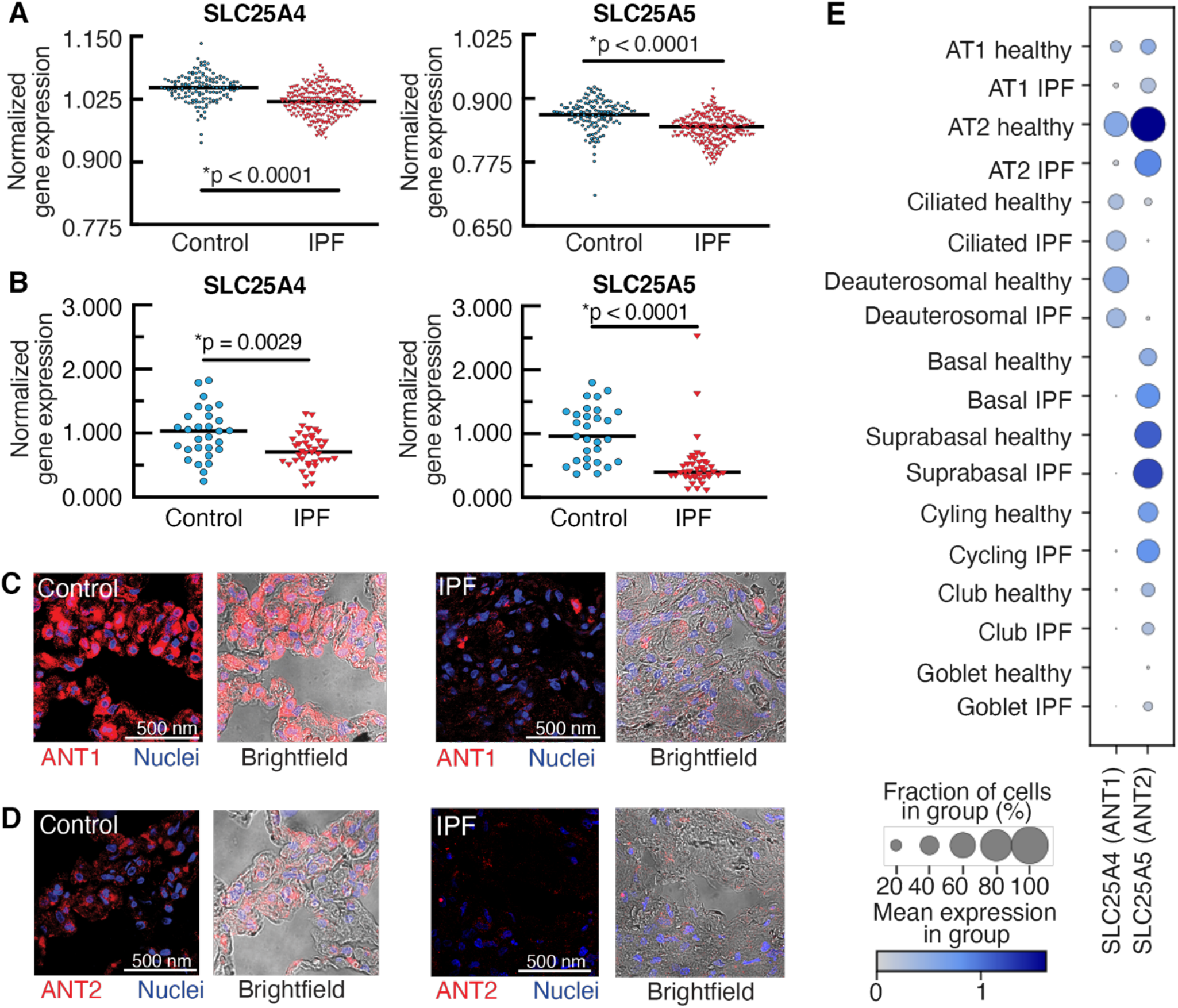
SLC25A4 (ANT1) and SLC25A5 (ANT2) expression is decreased in the lungs of patients with IPF. A) A human microarray dataset shows that gene expression of *SLC25A4* (*p < 0.0001) and *SLC25A5* (*p < 0.0001) is significantly decreased in whole lung lysates from IPF patients (n = 255) when compared to healthy controls (n = 137) (LGRC; 1RC2HL101715). Genomic data is normalized to glucose-6-phosphate isomerase (GPI). Statistics are by Student’s T test with Mann-Whitney post hoc test. Median bars are shown. B) RT-qPCR shows a significant decrease in *SLC25A4* (*P value = 0.0029) and *SLC25A5* (*p < 0.0001) gene expression in lung homogenates from a separate cohort of IPF patients (n = 40) compared to healthy control lungs (n = 29). Statistics are by unpaired nonparametric Mann-Whitney test. C,D) Representative immunofluorescence staining for ANT1 (Abcam #ab102032 rabbit polyclonal, 1:500), and ANT2 (Abcam #192410 rabbit polyclonal, 1:100) in human lung tissue sections from healthy controls and IPF patients. Images from additional patient samples are available in **Supplemental Figure 1** (n=4 subjects per group). Images were obtained by confocal microscopy at 60x. Brightfield overlay images are provided. Scale bar is 500 nm. E) Single cell RNA sequencing results obtained on isolated lung epithelial cells from IPF patients and normal controls. Mean expression levels and fraction of cells with gene expression of *SLC25*A4 and *SLC25A5* for each cell cluster type are depicted.

### Loss of ANT1 is associated with cellular senescence

Cellular senescence is associated with pro-aging stressors and increasingly implicated as a key mechanism in IPF pathogenesis (5). The cyclin-dependent kinase inhibitors p16^Ink4a^, p21^Cip1/Waf1^, p53, and positive beta-galactosidase staining are established markers of senescence (25,26). This notion that senescence is associated with IPF pathogenesis is supported in our LGRC whole lung tissue gene expression, which shows significant upregulation in *CDKN2A* (p16, *p < 0.0001) and *TP53* (p53, *p < 0.0001) gene expression in the lung tissue of the IPF cohort (**Figure 2A)**. Interestingly, we found that *CDKN2A* expression is inversely correlated to *SLC25A4* (ANT1) (*P value <0.0001) and *SLC25A5* (ANT2) (*P value <0.0001) expression in the IPF cohort (**Figure 2B)**, as well as in our single cell RNA sequencing data set from isolated lung epithelial cells from IPF lung tissue. Both *SLC25A4* (ANT1) and *SLC25A5* (ANT2) gene expression were inversely correlated to a number of common cellular senescence genes when analyzing all epithelial cells (**Figure 2C**) and specifically AT1 and AT2 cells (**Figure 2D**). Gene expression correlation tables are available in supplemental Tables E1 and E2. Together these data support that gene expression of *SLC25A4* (ANT1) and *SLC25A5* (ANT2) are reduced in epithelial cells in IPF and may influence cellular senescence in IPF.

**Figure 2:**
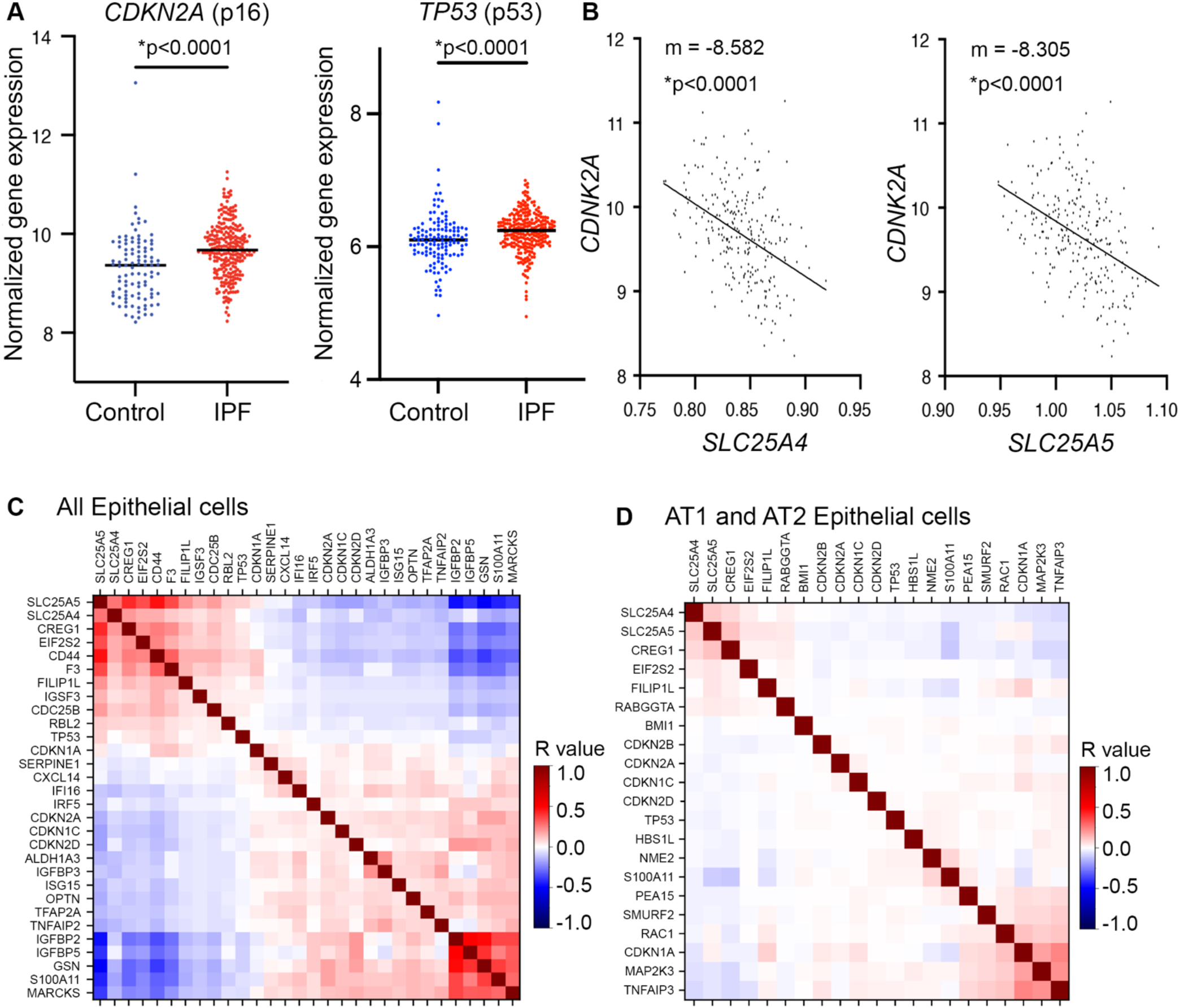
ANT gene expression inversely correlates with markers of cellular senescence. A) Gene expression data from the LGRC microarray dataset shows that *CDKN2A* (p16) (*p < 0.0001) and *TP53* (p53) are significantly increased in whole lung lysates from IPF patients (n = 255) when compared to lysates from healthy controls (n = 137). Statistics by a Students t test. P values are shown. B) Gene expression correlations between *CDKN2A* and *SLC25A4* (ANT1) (*P value < 0.0001) and *CDKN2A* and *SLC25A5* (ANT2) (*P value < 0.0001). Data is normalized to glucose-6-phosphate isomerase (GPI). Statistics are by Mann-Whitney unpaired nonparametric tests. Median bars are shown. C-D) Gene expression correlations were determined in the human lung epithelial single cell RNA sequencing data set for *SLC25A4* (ANT1), *SLC25A5* (ANT2) and common genes in the cellular senescence pathway for C) all epithelial cells or D) AT1 and AT2 epithelial cells. Color bar shows the calculated R value for each correlation.

To assess if loss of ANT1 expression influences the development of cellular senescence, we used small interfering RNA (siRNA) to suppress ANT1 expression in three immortalized human lung epithelial cell lines, then probed for markers of cellular senescence. Gene knockdown was confirmed using RT-qPCR (**Supplemental Figure E2**). In BEAS2B cells, ANT1 knockdown was associated with a significant increase in p21 (p value = 0.0220) and p53 (p value = 0.0205) by Western blot (**Figure 3A,B**). A trending but insignificant increase in p16 (P value = 0.0817) and p21 (P value = 0.1034) was observed in the HBEC3-KT cell line (**Figure 3C**). However, significantly (*P value = 0.0106) more HBEC3-KT cells stained positive for senescence associated beta-galactosidase stain and increased cell-body size with ANT1 knockdown compared to control treated cells (**Figure 3D**). Expression for p21 was also significantly increased in the alveolar epithelial cell line A549 with knockdown of ANT1 at baseline (**Figure 3E**). Next, we treated A549 cells with bleomycin to induce cellular senescence. Expression of p21 significantly increased in both the control (*P value <0.0001) and siANT1 (*P value <0.0001) groups with bleomycin treatment (**Figure 3E**) with a greater increase in p21 observed for the siANT1 knockdown group compared to controls (*P value = 0.0063). Similar trends were observed at the mRNA level; p21 significantly increased in both the control (*P value <0.0001) and siANT1 (*P value <0.0001) groups with bleomycin treatment (**Figure 3F**). These observations indicate that loss of ANT1 leads to increased cellular senescence at baseline and increased vulnerability to senescence stimuli in lung epithelial cells.

**Figure 3:**
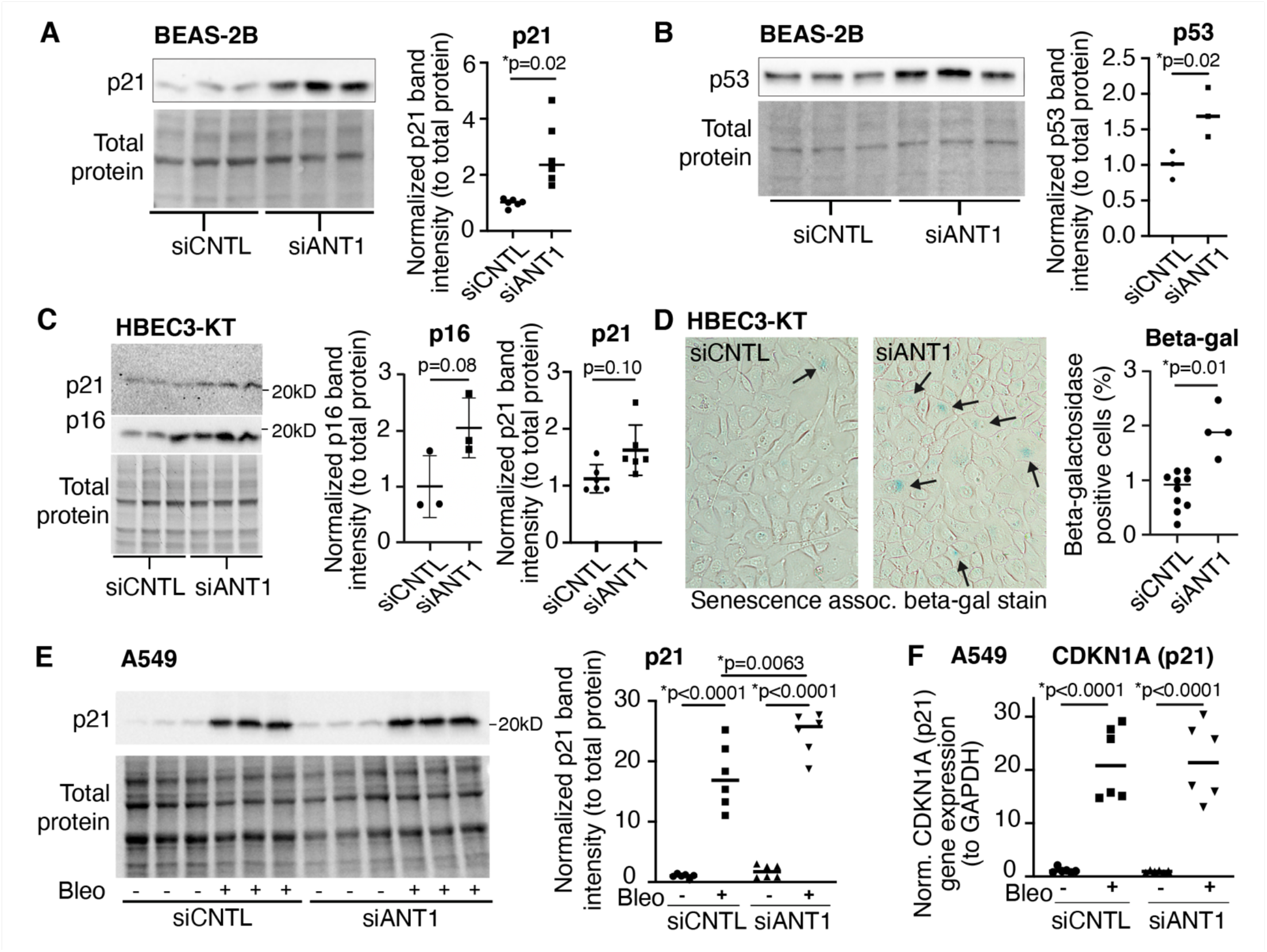
Loss of ANT1 results in increased markers of cellular senescence in multiple lung epithelial cell types. ANT1 expression was knocked down using siRNA compared to scrambled siRNA control conditions. Western blot and quantification are shown from human bronchial epithelial BEAS-2B cell lysates for A) p21 (*P value = 0.0220, n=6) and B) p53 (*P value = 0.0205, n=3). C) Western blot analysis was completed on human bronchial epithelial HBEC3-KT cells lysates show trending but insignificant increases in p16 (P value = 0.0817, n=3) and p21 (P value = 0.1034, n=6) with ANT1 knockdown. D) Increased senescence associated beta-galactosidase staining in HBEC3-KT cells with ANT1 knockdown compared to scrambled siRNA controls. Senescent cells are blue and noted by arrows. E) Western blot was completed on human alveolar A549 cells after ANT1 knockdown and probed for p21 (*P value = 0.0010, n=6). F) Gene expression for *CDKN1A* (p21) in A549 cells with ANT1 knockdown (*P value = 0.0063, n=6). All Western blot analysis band intensities are normalized to total protein intensity. Data shown are representative of 2-3 individual experiments. Statistics by unpaired t-test for comparisons between two groups, and by one-way ANOVA for comparing more than two groups. Error bars represent standard error.

### Loss of ANT1 results in worsened lung fibrosis and inflammation in mouse models of bleomycin and asbestos-injury

We next utilized a global Ant1 knock out (KO) mouse to determine how loss of Ant1 affects pulmonary fibrosis in the bleomycin and asbestos injury mouse model of pulmonary fibrosis. Gene knockdown was confirmed in heart, lung, kidney, and liver homogenates using RT-qPCR (**Supplemental Figure E3**). To model pulmonary fibrosis *in vivo*, mice were treated with bleomycin for 28 days. Notably, Ant1 KO mice treated with bleomycin have increased fibrosis and collagen deposition on H&E (**Figure 4A**) and trichrome staining (**Figure 4B**) compared to bleomycin-treated control mice. This is further supported by tissue scoring with a significant increase in fibrosis index in the bleomycin treated Ant1 KO mice (**Figure 4C**, *P value = 0.0255). There was no observed baseline lung phenotype in the Ant1 KO mice treated with saline compared to control mice. We next used intratracheal asbestos exposure as a second model of lung fibrosis. In the asbestos model, Ant1 KO mice treated with asbestos have increased fibrosis on H&E (**Supplemental Figure E4A**) and increased fibrosis index on tissue scoring (**Figure E4B**).

**Figure 4:**
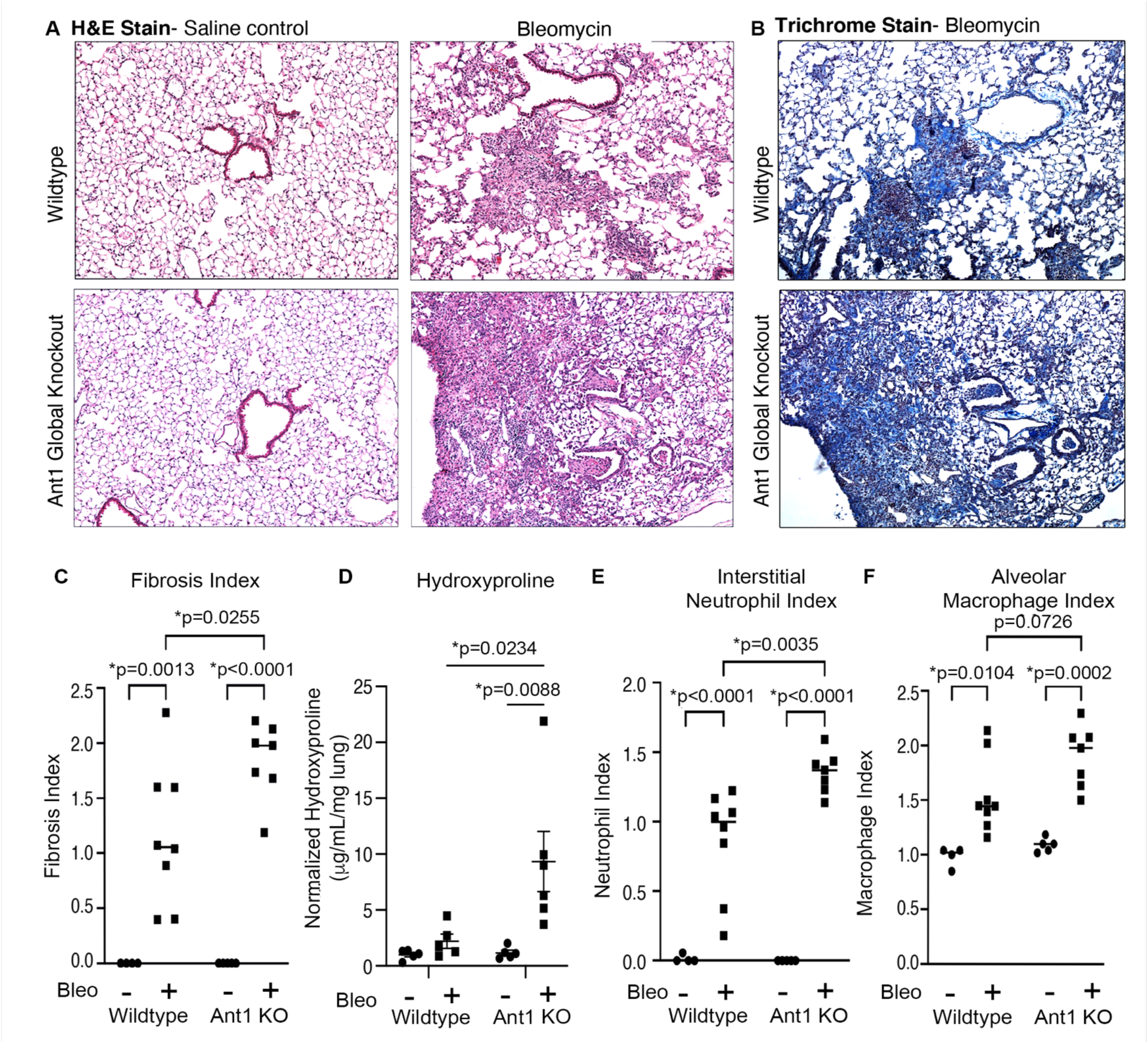
Global knock out of ANT1 leads to increased fibrosis and inflammation in murine lungs in a bleomycin model. A) Representative images from H&E staining of paraffin-embedded lung sections from Wildtype and Ant1 KO mice treated with saline or bleomycin for 28 days. B) Representative trichrome stain of paraffin-embedded lung sections from Wildtype and Ant1 KO mice treated with bleomycin for 28 days. C) Tissue scoring fibrosis index for fibrosis quantification. D) Hydroxyproline assay was completed on whole lung tissue samples (left lung lobe). Number of mouse lungs per group: WT saline (n = 5), WT bleomycin (n = 5), Ant1 KO saline (n = 5), Ant1 KO bleomycin (n = 6). Tissue scoring indices were determined for E) interstitial neutrophil index and F) alveolar macrophage Index. Statistics for all analyses are by ordinary One-Way ANOVA with Tukey post-hoc test.

Hydroxyproline is major component of fibrillar collagen and concentrations are directly correlated to total collagen in tissues. To further support our finding that loss of Ant1 results in increased lung fibrosis, we quantified hydroxyproline on the left lung lobe from the mice to assess collagen content. A significant increase in hydroxyproline concentration was observed in lungs from bleomycin-treated Ant1 KO mice compared to saline-treated controls (**Figure 4D**, *P value = 0.0088) and compared to bleomycin-treated WT mice (**Figure 4D**, *P value = 0.0234). There was a non-significant small trend towards increased hydroxyproline in WT mice between bleomycin and saline-treated groups.

Cellular senescence is associated with the secretion of a milieu of cytokines, chemokines, growth factors, matrix proteases, and other inflammatory factors, collectively known as the senescence associated secretory phenotype (SASP) secretome. SASP signaling has been proposed as a mechanism driving inflammation and tissue remodeling in the fibrotic lung (5). In order to quantify lung tissue inflammation, neutrophil and macrophage infiltration was scored from H&E-stained tissue sections. Bleomycin induced pulmonary fibrosis was associated with an increase in neutrophil and macrophage indices in both WT and Ant1 KO mice (**Figure 4E,F**). Loss of Ant1 is associated with a significantly greater increase of neutrophils (*P value = 0.0035) and a trending increase in macrophage infiltration (P value = 0.0726). In the asbestos model, there were no significant differences in neutrophil or macrophage inflammation indices between the WT and Ant1KO asbestos-treated groups (**Figure E4C,D**). Together, these data demonstrate that loss of Ant1 leads to increased lung fibrosis, collagen deposition and inflammation in the bleomycin and asbestos lung injury models.

### Loss of Ant1 results in increased markers of senescence and senescence associated secretory phenotype(SASP) in the lung

To probe for markers of cellular senescence *in vivo*, we stained mouse lung sections against p21 proteins (**Figure 5A**). We quantified nuclear levels of p21 staining within the alveolar regions of the lung tissue with Fiji (see Expanded Materials and Methods). We found no change in nuclear p21 staining between the saline groups. A significant increase in p21 staining was noted for the bleomycin treated Ant1KO mice (*P value < 0.0001) when compared to the saline groups and WT bleomycin (**Figure 5A,B**). Staining against p21 was notably higher in the bleomycin Ant1KO mice compared to the bleomycin-treated WT mice.

**Figure 5:**
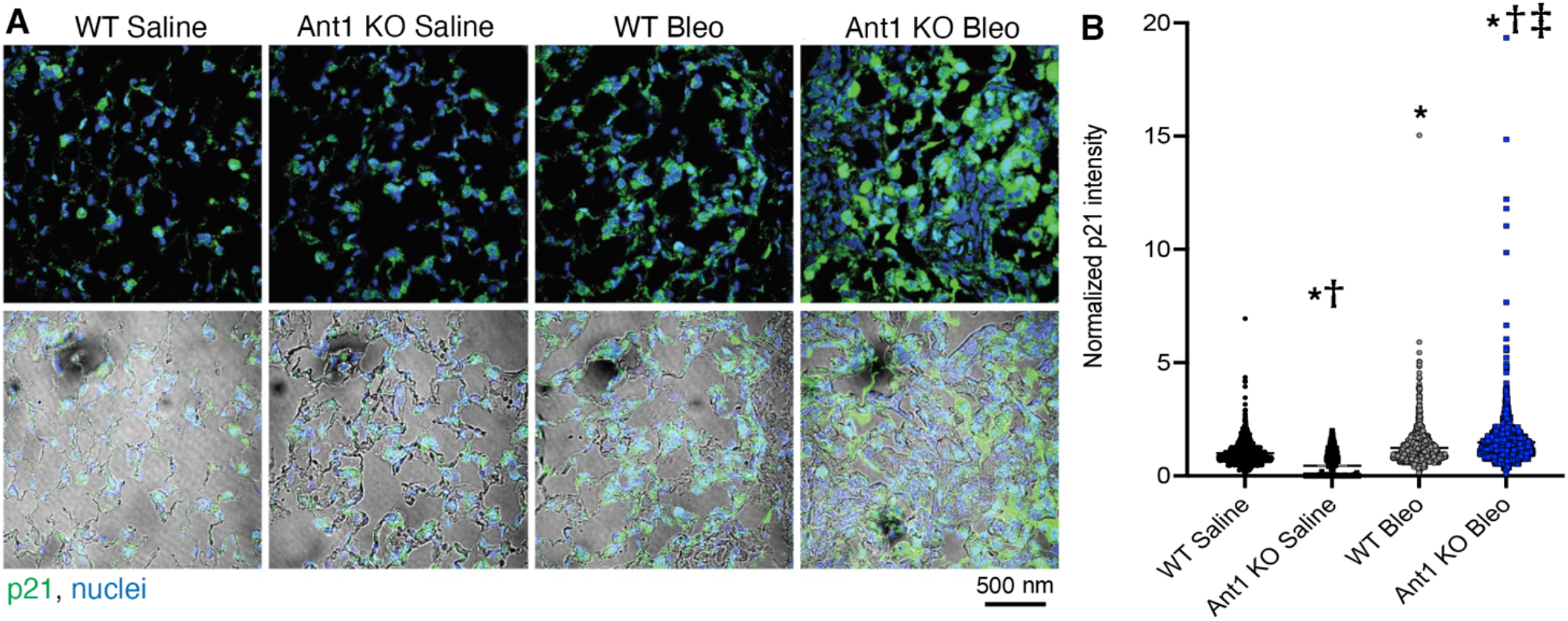
Loss of Ant1 results in increased p21 in pulmonary alveolar tissue after bleomycin injury. Wildtype (WT) and Ant1 KO mice were exposed to bleomycin (28 days) and lung tissue was stained for p21 using immunofluorescence (IF) with confocal imaging. A) Representative p21 staining of alveolar lung tissue. Green – p21, Blue – DAPI-stained nuclei. Bottom row of images shows a brightfield image overlay for lung structure. B) Quantification of nuclear p21 intensity in alveolar tissue areas. Statistics are by One-way ANOVA with Kruskal-Wallis. N = 10-18 images per mouse with 3 mice per group. P values are: *p<0.0001 versus WT saline; †p<0.0001 versus WT bleomycin; ‡p<0.0001 versus Ant1KO saline.

Senescence due to mitochondrial dysfunction has been shown to produce a unique SASP from other senescence stimuli (8) which may be tissue and injury specific. We found that several secreted growth factors and pro-inflammatory factors associated within the SASP secretome are significantly increased in the bronchioalveolar lavage (BAL) from bleomycin-treated Ant1 KO mice, but not WT mice, when compared to saline-treated controls (**Figure 6 and Table 1**, additional data in Supplemental Figures E5 and E6). We find that loss of Ant1 is associated with a significant enrichment in secreted matrix metalloproteinase 12 (MMP-12, *P value < 0.0001), growth/differentiation factor 15 (GDF-15, *P value = 0.0139), and vascular endothelial growth factor (VEGF, *P value = 0.0251) in BAL from bleomycin-treated mice compared to saline-treated controls. This data supports the finding that loss of Ant1 leads to enhanced tissue remodeling in the lung parenchyma following bleomycin treatment. A significant increase in proinflammatory chemokines was also observed in the BALs of Ant1 KO mice with bleomycin treatment including chemokine (C-C motif) ligand 3. This chemokine is a key driver of inflammatory cell recruitment, including monocytes and other leukocytes, to the lung environment. There was also an increase in additional proinflammatory markers including CCL8 (*P value = 0.0117) and granulocyte-macrophage colony stimulating factor (GM-CSF, *P value = 0.0465) in BALs from bleomycin-treated Ant1 KO mice. Additional factors associated with inflammation, are also enriched in BALs from bleomycin treated Ant1 KO mice. A trending but insignificant increase in CXCL1/GRO alpha (P value = 0.0883) and interleukin-7 (IL-7) is also observed in BALs from bleomycin treated Ant1 KO mice. A complete list summarizing significant changes in cytokine concentrations is available in **Table 1**. This data suggests that loss of Ant1 contributes to the increased pro-inflammatory response in the lung parenchyma following bleomycin insult by modulating the SASP secretome.

**Table 1:** Cytokines measured in bronchioalveolar lavage samples

| **SASP factor** | **Type of cytokine** | **WT mice,**<br>**Bleo vs saline** | **Ant1 KO mice,**<br>**Bleo vs saline** | **Ant1 KO vs WT**<br>**Bleomycin groups** |
| --- | --- | --- | --- | --- |
| MMP-12 | Proteases and regulators |  | Up with bleo,<br>*P value<br><0.0001 | Up with Ant1 KO,<br>*P value = 0.0007 |
| MMP-3 | Proteases and regulators | Up with bleo<sup>†</sup> | Up with bleo (P<br>value<br>unavailable) | Undetermined<sup>†</sup> |
| ICAM-1/CD54 | Soluble or shed receptors<br>or ligands |  | Up with bleo,<br>*P value =<br>0.0022 |  |
| FasL/TNFSF6/<br>CD95L/CD178 | Soluble or shed receptors<br>or ligands |  | Trending up<br>with bleo, *P<br>value = 0.0823 |  |
| GDF-15/MIC-1 | Growth factor or regulator |  | Up with bleo,<br>*P value =<br>0.0139 | Trending up with<br>Ant1 KO, *P value<br>= 0.1694 |
| VEGF | Growth factor or regulator |  | Up with bleo,<br>*P value =<br>0.0251 |  |
| IGFBP-3 | Growth factor or regulator | Up with bleo, *P<br>value = 0.0090 | Up with bleo,<br>*P value =<br>0.0343 |  |
| Osteopontin/OPN | Growth factor or regulator | Up with bleo<sup>†</sup> | Up with bleo,<br>*P value =<br>0.0405 | Undetermined<sup>†</sup> |
| CCL4/MIP-1 beta | Chemokine | Trending up with<br>bleo, P value =<br>0.1269 | Up with bleo,<br>**P value =<br>0.0010 |  |
| CCL3/MIP-1<br>alpha | Chemokine |  | Up with bleo,<br>*P value =<br>0.0073 |  |
| CLC8/MCP-2 | Chemokine |  | Up with bleo,<br>*P value =<br>0.0117 |  |
| CXCL2/GRO<br>beta | Chemokine |  | Up with bleo,<br>*P value =<br>0.0179 |  |
| CCL22/MDC | Chemokine |  | Up with bleo,<br>*P value =<br>0.0407 |  |
| CXCL1/GRO<br>alpha | Chemokine |  | Trending up<br>with bleo, P<br>value = 0.0883 |  |
| IL-1 beta | Interleukin | Up with bleo, *P<br>value = 0.0482 | Up with bleo,<br>*P value =<br>0.0499 |  |
| IL-7 | Interleukin |  | Trending up with bleo, P value = 0.0622 |  |
| GM-CSF | Other inflammatory factor |  | Up with bleo, *P value = 0.0465 |  |
| M-CSF | Other inflammatory factor |  | Trending up with bleo, P value = 0.0883 |  |

Not shown: No significant change was observed between groups for chemokines CCL7; CCL2/MCP-1; LIX/CXCL5/ENA-78, interleukins IL-17; IL-1 alpha, and other inflammatory factor IFN-gamma. Blank cells indicate no significant or trending changes in cytokine concentrations between groups.
† The cytokine concentrations from the bleomycin group were measured above detection range and therefore not available to calculate P value.

**Figure 6:**
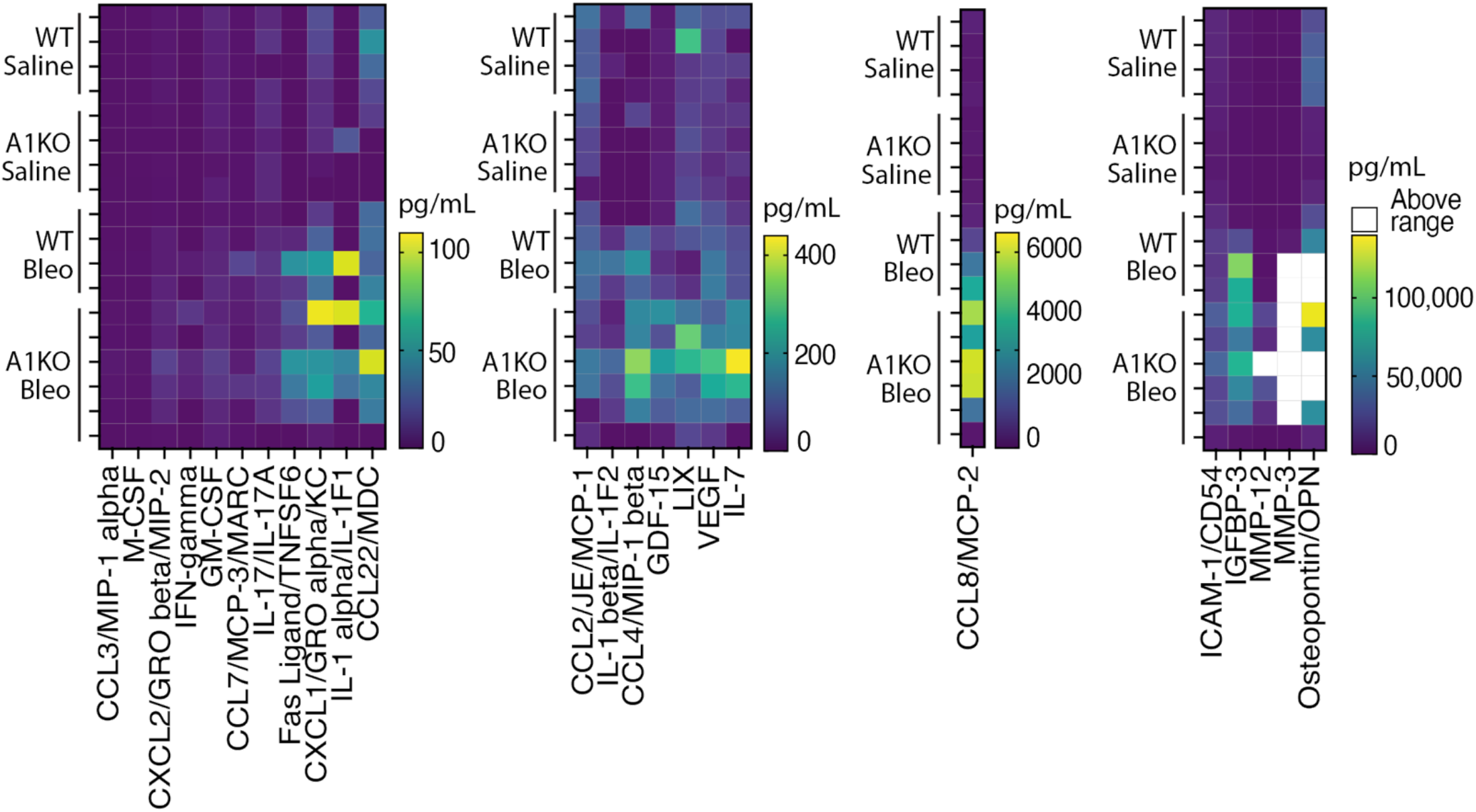
Increased SASP identified in the bronchioalveolar lavage with loss of Ant1 and bleomycin injury. Heat maps representing cytokine concentrations in bronchoalveolar lavage of WT saline (n=3), Ant1 KO saline (n=4), WT bleomycin (n=4), and Ant1 KO bleomycin (n=6) at 28 days post-treatment. Cytokine concentrations are shown in pg/mL by color gradients. Select concentrations were above the detection range (white).

## Discussion

Cellular senescence and mitochondrial dysfunction have been implicated in the pathogenesis of IPF, however the factors important for driving mitochondrial dysfunction-associated senescence (MiDAS) remain unclear. The present study is the first to demonstrate that ANT1, a mitochondrial ATP/ADP antiporter, is critical for the regulation of cellular senescence in the lung and IPF. Markers of senescence have been identified in epithelial cells within the lungs of patients with IPF (9). In IPF, AT2 alveolar epithelial cells have altered mitochondrial function represented by a reduction in NAD/NADH ratio, reduced ATP and decreases in the ETC complexes (21). Leveraging human IPF lung tissue, we determine that ANT1 and ANT2 are reduced in the AT1 and AT2 lung epithelial cells of patients with IPF, suggesting that depletion of ANTs and subsequent mitochondrial dysfunction are implicated in IPF pathogenesis.

ANTs are abundant ATP/ADP transporters across the inner mitochondrial membrane and are integral to maintaining energy balance in the cell. We have previously demonstrated a direct relationship between loss of ANT1 expression and cellular energetics in lung epithelial cells, such as decreases in mitochondrial ATP production and basal oxygen consumption rate (21). Mitochondrial dysfunction in IPF has been identified by decreased activity in the electron transport chain (ETC), reduced ATP production (27) and increased production of mitochondrial reactive oxygen species (ROS) (28–31). Mitochondrial dysfunction exacerbates alveolar cell injury and may contribute to fibrogenesis (32–34). Here, we have shown that loss of ANT1 *in vitro* in cultured human epithelial lung cells and *in vivo* loss of ANT1 results increased markers of senescence, namely p16^Ink4a^, p21^Cip1/Waf1^, p53, and positive beta-galactosidase. Furthermore, in two mouse models of lung fibrosis, Ant1 deletion resulted in worsened lung fibrosis through enhanced cellular senescence in the lung epithelium. These mice express increased SASP markers in BAL fluid, including chemokines and growth factors such as GDF-15, a biomarker of epithelial stress upregulated in lungs of IPF patients (35). This increase in secreted SASP factors likely contributes to the propagation of lung injury seen in the lung with loss of Ant1. These findings highlight the importance of understanding how SASP may change with different methods of MiDAS induction. Prior studies of MiDAS utilized depletion of mitochondrial DNA (8) to induce mitochondrial dysfunction and stress with a unique SASP that lacked IL1 signaling. In our study we found intact IL-1 response but enhanced growth factors and chemokines, suggesting that MiDAS SASP induced by loss of ANTs in the context of injury differs and requires further investigation.

While our studies indicate that global loss of Ant1 in transgenic mice predispose epithelial cells to become senescent and causes worse fibrosis in mouse fibrosis models, one limitation is that Ant1 is depleted in all cells in this global knockout mouse model. Our data suggests that epithelial cells are involved in the induction of cellular senescence due to loss of ANT1, however It is unclear if other cell types (such as immune cells) play a role which may indirectly contribute to this process. Future studies could elucidate this using cell-specific alterations of Ant1 expression. For *in vitro* studies, given that alveolar A549 cells are an adenocarcinoma-based cell, we also performed these in bronchial epithelial cells lines that are not cancer derived, with the limitation that bronchial cells may behave differently than alveolar epithelial cells.

Loss of ANTs result in a reduction in mitochondrial oxidative respiration and ATP production (21) due to less transport of ATP out of mitochondria, resulting in a backup of the electron transport chain. Emerging evidence suggests that sensors of the energetic and metabolic state of the cell are important in senescence, specifically NAD+/NADH ratios and AMPK (8,36). Whether depletion of ANTs alters these metabolic sensors to drive cellular senescence remains to be elucidated by future studies. Our findings establish that pulmonary fibrosis is in part mediated by ANT-related MiDAS and provides a potential therapeutic target for the treatment of IPF to limit fibrosis development. ANTs have been proposed as a possible method of cell-specific targeting of senolytic drugs, as previous studies report that ANT-deficient cells are more susceptible to the cytotoxic effect of the drug MitoTam (37).

In conclusion, our findings suggest that ANTs may be key drivers of MiDAS in the lung and present a potential therapeutic target for limiting fibrosis development in IPF. Therapeutic opportunities could involve senolytics or modulation of ANT expression to tune metabolic state and function in epithelial cells. The findings of this study underscore the importance of understanding the role of ANT proteins in senescence, MiDAS and lung disease.

## Supporting information

Supplemental - Supp_tableE1_AllEpithCells_SLC25A4_SLC25A5_correlations_IPF

Supplemental - Supp_tableE2_AT2_AT1 cells_SLC25A4_SLC25A5_correlations_IPF.xls

## Abbreviations

A549: adenocarcinoma alveolar epithelial cell line
ADP: adenosine diphosphate
ATP: adenosine triphosphate
ANT1: Adenine Nucleotide Translocase 1
ANT2: Adenine Nucleotide Translocase 2
AT1: Alveolar type 1 cell
AT2: Alveolar type 2 cell
BCA: bicinchoninic acid
BEAS2B: human bronchial epithelial cell line
β-gal: beta-galactosidase stain
BSA: bovine serum albumin
CCL2/JE/MCP-1: CC Motif Chemokine Ligand 2/Monocyte-Chemotactic Protein 1
CCL3/MIP-1 alpha: CC Motif Chemokine Ligand 3/Macrophage Inflammatory Protein-1 alpha
CCL4/MIP-1 beta: CC Motif Chemokine Ligand 4/Macrophage Inflammatory Protein-1 beta
CCL7/MCP-3/MARC: CC Motif Chemokine Ligand 7/Monocyte-Chemotactic Protein 3
CCL8/MCP-2: CC Motif Chemokine Ligand 8/Monocyte Chemotactic Protein-2
CCL22/MDC: CC Motif Chemokine Ligand 22/Macrophage-derived Chemokine
CXCL1/GRO Alpha/KC/CINC-1: CXC Motif Chemokine Ligand 1/Growth-Regulated Oncogene alpha
CXCL2/GRO Beta/MIP-2/CINC-3: CXC Motif Chemokine Ligand 2/Growth-Regulated Oncogene beta
CXCL5/LIX: CXC Motif Chemokine Ligand 5
DMEM: Dublecco’s modified Eagle’s medium
DMSO: dimethyl sulfoxide
DNA: deoxyribonucleic acid
DNase: deoxyribonuclease
DTT: dithiothreitol
EDTA: ethylenediaminetetraacetic acid
ETC: Electron transport chain
FasL/TNFSF6: Fas-Ligand
FBS: fetal bovine serum
FITC: fluorescein isothiocyante
g: gram
GAPDH: glyceraldehye 3-phospate dehydrogenase
GDF-15: Growth/Differentiation Factor 15
GPI: glucose-6-phosphate isomerase
GM-CSF: Granulocyte-macrophage Colony Stimulating Factor
GWAS: genome-wide association study
h: hour
H&E: hematoxylin and eosin
HBEC3-KT: human bronchial epithelial cell line
IF: immunofluorescence
(IGFBP-3): Insulin-like Growth Factor Binding Protein 3
IPF: Idiopathic Pulmonary Fibrosis
ICAM-1/CD54: Intercellular Adhesion Molecule 1
IL-1 alpha/IL-1F1: Interleukin 1 alpha
IL-1 beta/IL-1F2: Interleukin 1 beta 1
IL-7: Interleukin 7
IL-17/IL-17A: Interleukin 17
IFN-gamma: Interferon gamma
IT: intratracheal
kD: kilodalton(s)
LGRC: lung genome research consortium
M-CSF: Macrophage Colony Stimulating Factor
MiDaS: Mitochondrial dysfunction associated cellular senescence
MMP-3: Matrix Metalloproteinase 3
MMP-12: Matrix Metalloproteinase 12
mL: milliliter(s)
μL: microliter(s)
μm: micrometer(s)
OCR: oxygen consumption rate
OPN: Osteopontin
PMN: polymorphic neutrophils
RIPA buffer: radioimmunoprecipitation assay buffer
RNA: ribonucleic acid
ROI: region of interest
ROS: reactive oxygen species
scRNAseq: single cell RNA sequencing
SDS: sodium dodecyl sulfate
SEM: standard error of the mean
SLC25A4: gene for Adenine Nucleotide Translocase 1
SLC25A5: gene for Adenine Nucleotide Translocase 2
t test: Student’s t test
TRITC: tetramethyl rhodamine isothiocyante
V: volt
VEGF: Vascular Endothelial Growth Factor
vol: volume
RT-qPCR: real-time quantitative polymerase chain reaction
W: watt
wk: week
wt: weight
yr: year

## Acknowledgements

We would like to thank Mary Jane Duermeyer, Christine Burton, Ian Walsh, Jian Shi, Yadong Xiao and Yanwu Zhao for their technical assistance.

