## Supplemental figures and methods for "Loss of ANT1 Increases Fibrosis and Epithelial Cell Senescence in Idiopathic Pulmonary Fibrosis"

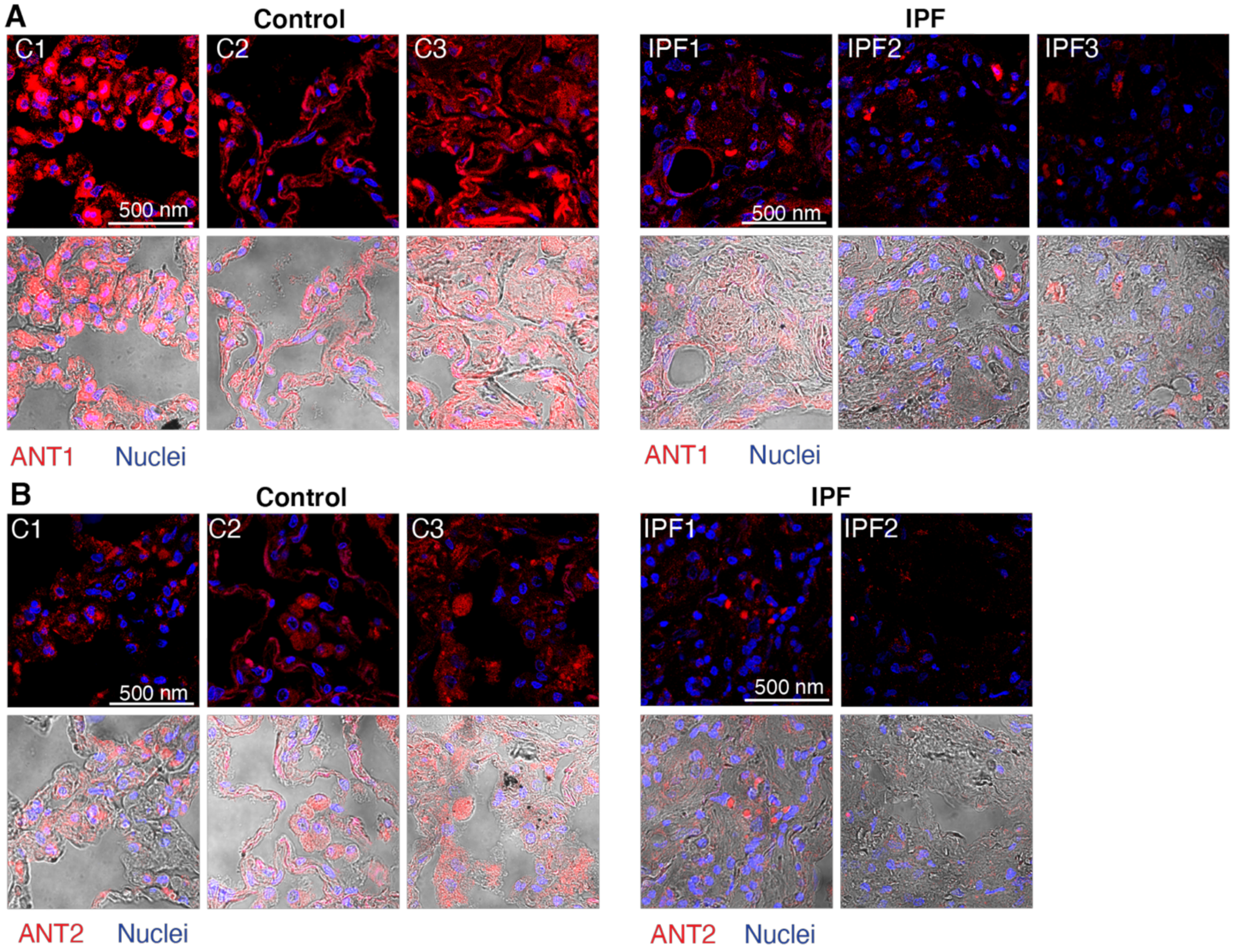


**Supplemental Figure E1: ANT1 and ANT2 staining is decreased in human lungs tissue from patients with IPF patients compared to control lung.**

Additional representative IF staining of human lung tissue sections from healthy controls (n=3) and IPF patients (n=3) for **A)** ANT1 (Abcam #ab102032 rabbit polyclonal, 1:500), and **B)** ANT2 (Abcam #192410 rabbit polyclonal, 1:100) displayed in red (dapi nuclei are in blue). Images obtained by confocal microscopy with a 60x objective. Brightfield overlay images are provided. Scale bar is 500 nm.

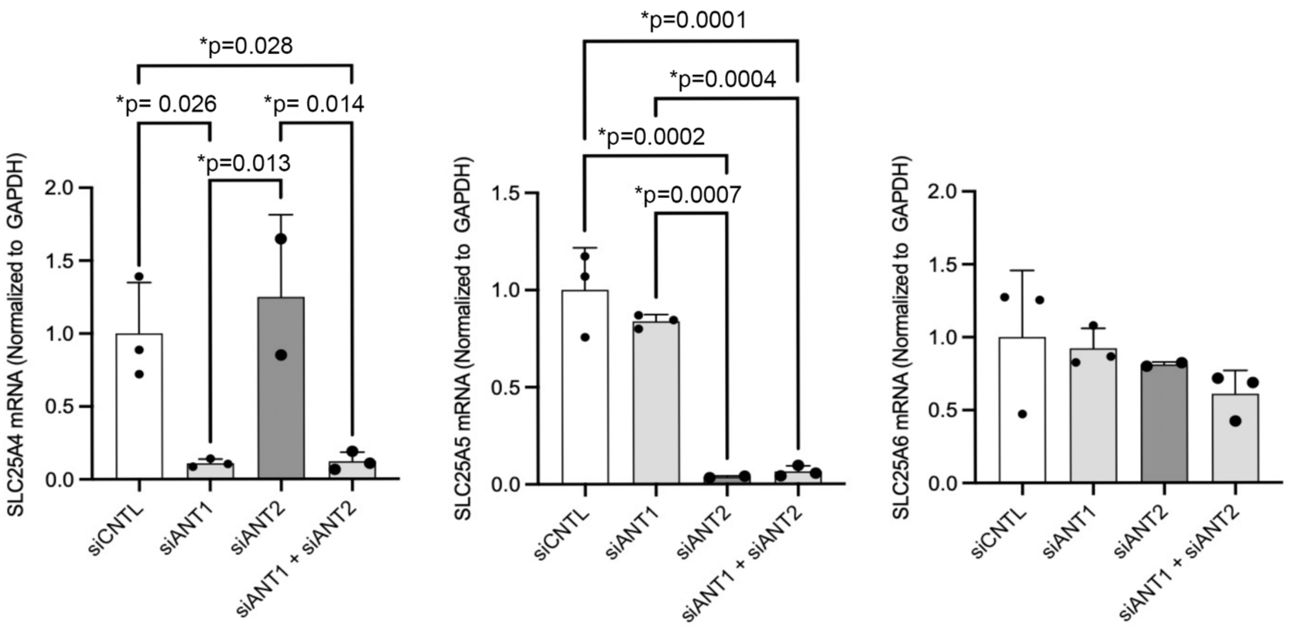


**Supplemental Figure E2: Confirmation of ANT gene expression knockdown in human bronchial epithelial cells, BEAS-2Bs.** siRNA was utilized to knockdown *SLC25A4* (siANT1) and/or *SLC25A5* (siANT2) in Beas-2b cells with scrambled sequence siRNA used as control (siCNTL). Gene expression knockdown was confirmed by RT-qPCR for **A)** *SLC25A4* (ANT1), **B)** *SLC25A5* (ANT2),and **C)** *SLC25A6* (ANT3, expressed only in human cells). Statistics by one-way ANOVA with Tukey's post-hoc test. P values are noted.

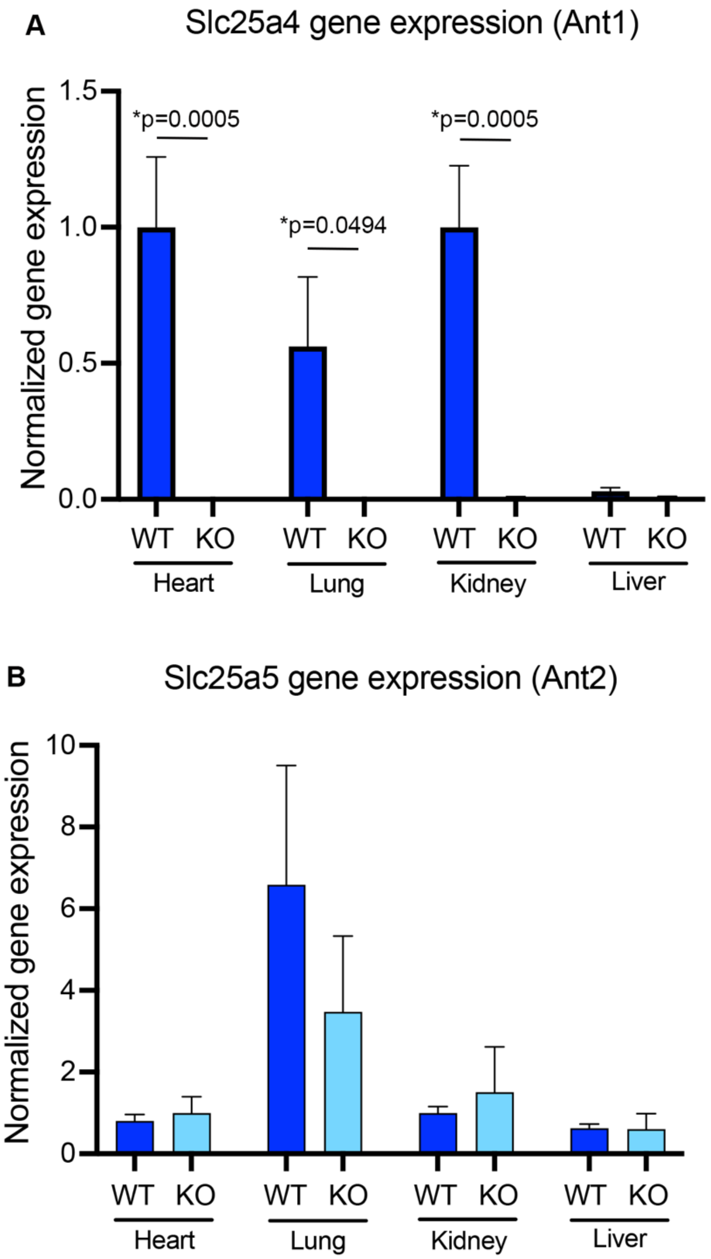


**Supplemental Figure E3: Confirmation of the Ant1 knock out mouse.** RT-qPCR was used to determine gene expression for ANT genes in mouse tissue including heart, lung, kidney and liver from WT and Ant1 KO mice (n=3 mice per group). **A)** *SLC25A4* **(**Ant1) and **B)** *SLC25A5* **(**Ant2) gene expression. Statistics by Two-way ANOVA with Bonferroni post-test.

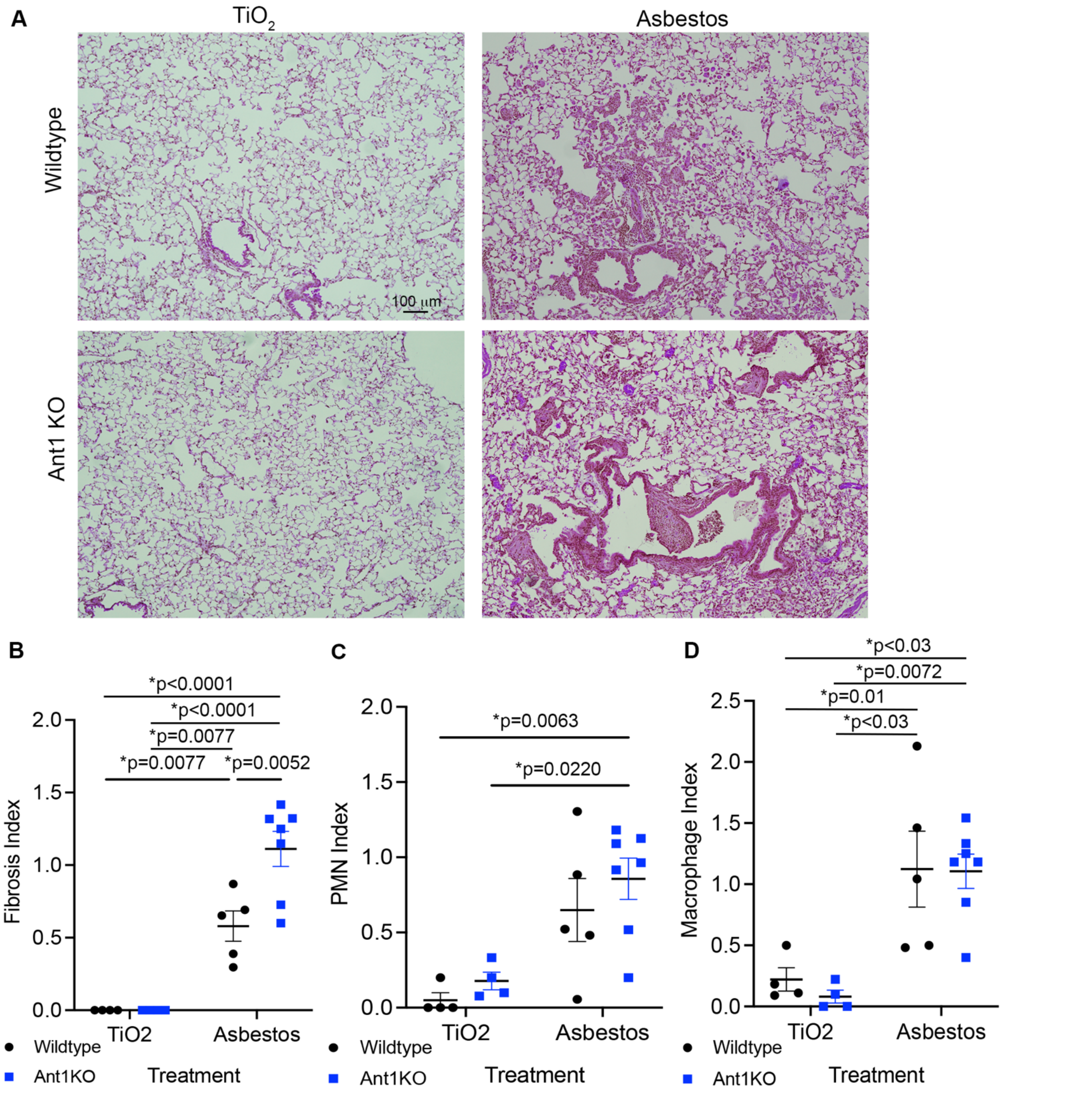


**Supplemental Figure E4: Global knock out of ANT1 leads to increased fibrosis and inflammation in murine lungs in an asbestos fibrosis model.** A) Representative images from H&E staining of paraffin-embedded lung sections from Wildtype and Ant1 KO mice treated with titanium dioxide control ($TiO_2$) or crocidolite asbestos for 28 days, n=4-7 mice per group. B-D) Tissue scoring indices for C) fibrosis index, D) interstitial neutrophil (PMN) index, and E) alveolar macrophage Index. Statistics by Two-Way ANOVA with Tukey post-hoc test. P values are noted.

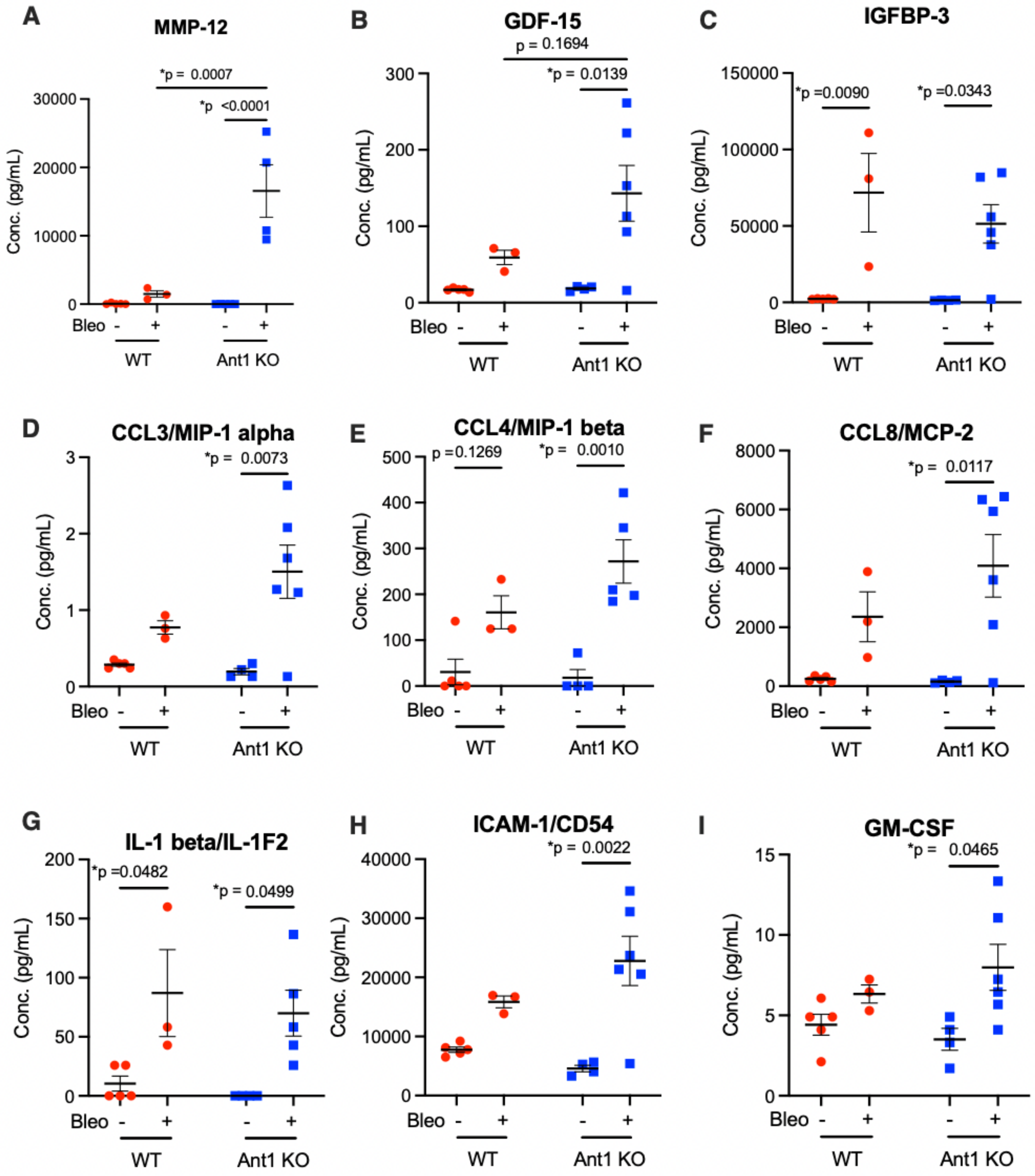


**Supplemental Figure E5: Increased SASP markers in bronchioalveolar lavage with bleomycin treatment in Ant1 KO mice.** Cytokine concentrations in bronchoalveolar lavage from WT and Ant1 KO mice treated with bleomycin form 28 days. A significant increase in analyte concentrations are noted in Ant1 KO mice treated with bleomycin: A) Matrix Metalloproteinase 12 (MMP-12, *P value < 0.0001); B) Growth/Differentiation Factor 15 (GDF-15, *P value = 0.0139); C) Insulin-like Growth Factor Binding Protein 3(IGFBP-3, *P value = 0.0343); D) CC Motif Chemokine Ligand 3/Macrophage Inflammatory Protein-1 alpha (CCL3/MIP-1 alpha, *P value =

0.0073); E) CC Motif Chemokine Ligand 4/Macrophage Inflammatory Protein-1 beta (CCL4/MIP-1 beta, *P value = 0.0010); F) CC Motif Chemokine Ligand 8/Monocyte Chemotactic Protein-2 (CCL8/MCP-2, *P value = 0.01117); G) Interleukin 1 beta 1 (IL-1 beta/IL-1F2, *P value = 0.0499); H) Intercellular Adhesion Molecule 1 (ICAM-1/CD54, *P value = 0.0022); I) Granulocyte-macrophage Colony Stimulating Factor (GM-CSF, *P value = 0.0465). Statistics are by 2-way ANOVA with Tukey post-hoc test. Bars represent median values. Number of mice per group: WT saline (n = 5), WT bleomycin (n = 3), Ant1 KO saline (n = 4), Ant1 KO bleomycin (n = 6). For MMP-12, only 4 values are shown from the Ant1 KO bleomycin group; 2 additional values were measured but were above detection range. Additional data is available in **Supplemental Figure E6** and **Table 1**.

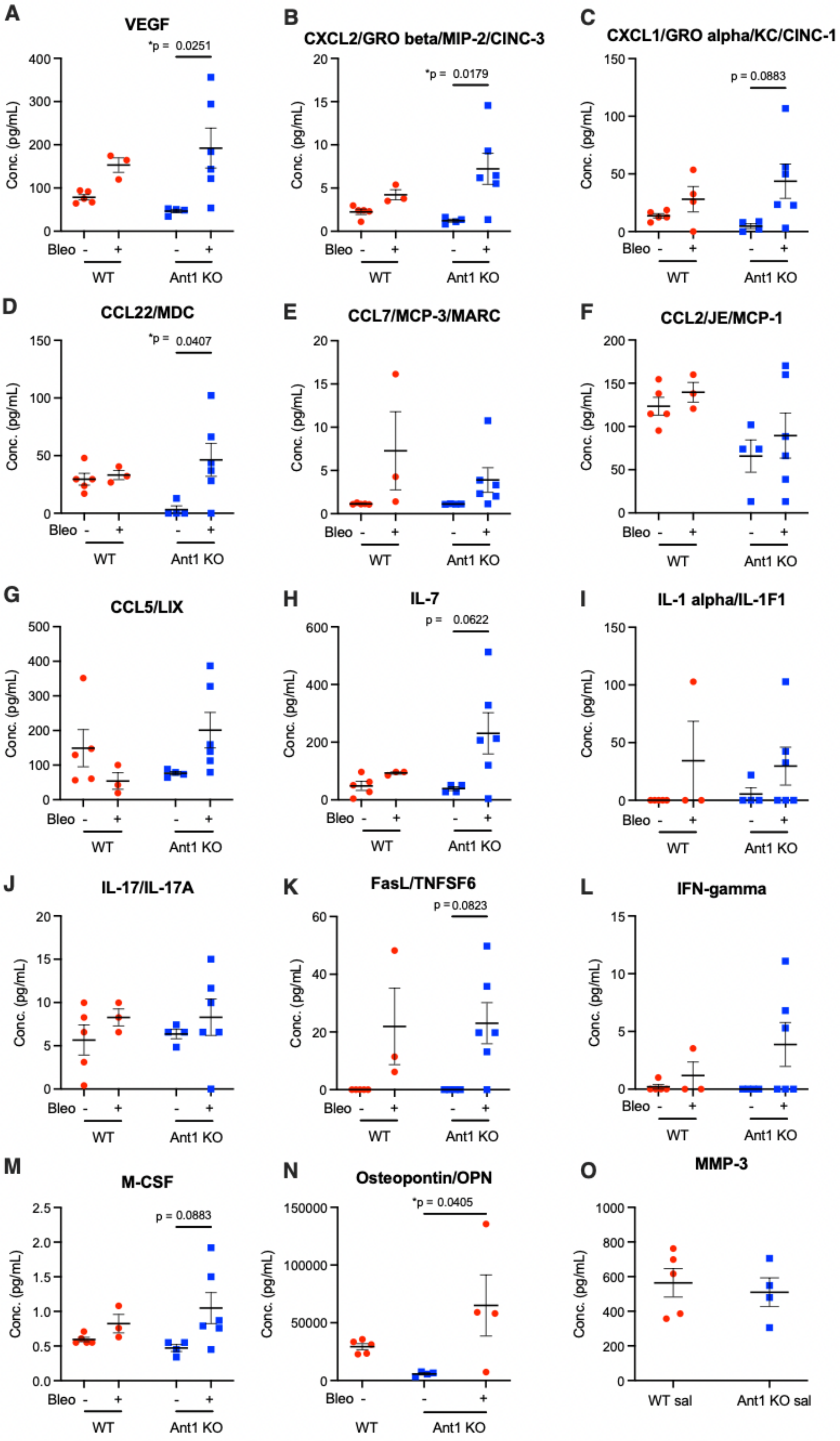
A
VEGF
*p = 0.0251
Conc. (pg/mL)
Bleo - + - +
WT
Ant1 KO
B
CXCL2/GRO beta/MIP-2/CINC-3
*p = 0.0179
C
CXCL1/GRO alpha/KC/CINC-1
p = 0.0883
D
CCL22/MDC
*p = 0.0407
E
CCL7/MCP-3/MARC
F
CCL2/JE/MCP-1
G
CCL5/LIX
H
IL-7
p = 0.0622
I
IL-1 alpha/IL-1F1
J
IL-17/IL-17A
K
FasL/TNFSF6
p = 0.0823
L
IFN-gamma
M
M-CSF
p = 0.0883
N
Osteopontin/OPN
*p = 0.0405
O
MMP-3
WT sal
Ant1 KO sal

**Supplemental Figure E6: Cytokine levels in bronchioalveolar lavage with bleomycin treatment in Ant1 KO mice.** Cytokine concentrations in bronchioalveolar lavage from bleomycin and saline treated Ant1 KO and Wildtype mice for A) Vascular Endothelial Growth Factor (VEGF); B) CXC Motif Chemokine Ligand 2/Growth-Regulated Oncogene beta (CXCL2/GRO Beta/MIP-2/CINC-3); C) CXC Motif Chemokine Ligand 1/Growth-Regulard Oncogene apha (CXCL1/GRO Alpha/KC/CINC-1); D) CC Motif Chemokine Ligand 22/Macrophage-derived Chemokine (CCL22/MDC); E) CC Motif Chemokine Ligand 7/Monocyte-Chemotactic Protein 3 (CCL7/MCP-3/MARC); F) CC Motif Chemokine Ligand 2/Monocyte-Chemotactic Protein 1 (CCL2/JE/MCP-1); G) CXC Motif Chemokine Ligand 5 (CXCL5/LIX); H) Interleukin 7 (IL-7); I) Interleukin 1 alpha (IL-1 alpha/IL-1F1); J) Interleukin 17 (IL-17/IL-17A); K) Fas-Ligand (FasL/TNFSF6); L) Interferon gamma (IFN-gamma); M) Macrophage Colony Stimulating Factor (M-CSF); N) Osteopontin (OPN); O) Matrix Metalloproteinase 3 (MMP-3);. Additional graphs and information are available in **Figure E5** and **Table 1**. Statistics are by 2-way ANOVA with Tukey post-hoc test. Bars represent median values. Number of mice per group: WT saline (n = 5), WT bleomycin (n = 3), Ant1 KO saline (n = 4), Ant1 KO bleomycin (n = 6).

**Supplemental Tables E1 and E2**

**Correlation tables** for cellular senescence genes correlated to SLC25A4 (ANT1) and SLC25A5 (ANT2) using the single cell RNA sequencing data set from epithelial cells isolated from IPF lung tissue representing either “all epithelial cells” (**Table E1**) or “AT1 and AT2” cells (**Table E2**). See the expanded methods for additional details.

**Supplemental Expanded Materials and Methods**

*Human lung tissue studies*

Human lung tissue samples were obtained from The Airway Cell and Tissue Core at the University of Pittsburgh (supported by P30 DK072506, NIDDK and the CFF RDP to the University of Pittsburgh). Donor lung samples were obtained from the Center for Organ Recovery and Education (CORE) at the University of Pittsburgh. Donor lung samples originated from lungs deemed unsuitable for organ transplantation. All IPF samples were from lungs explanted from IPF patients that had undergone lung transplantation under an approved protocol (STUDY18100070). Genomic data was obtained from the Lung Genomics Research Consortium (LGRC) (GEO GSE47460; http://www.lung-genomics.org/) using tissue samples and clinical data collected through the Lung Tissue Research Consortium (LTRC; http://www.ltrcpublic.com/). Lung tissues were stored at -80 °C until used. Lung homogenates were prepared in a Bullet Blender, and ANT expression levels were measured by Western analysis and RT-qPCR (see below). RT-qPCR for *SLC25A4* and *SLC25A5* was performed on mRNA extracted from lung tissue samples from control individuals (n=29) and IPF patients (n=40) and was normalized to glyceraldehye 3-phospate dehydrogenase (GAPDH).

*Immunocytochemistry of mouse and human lung tissue sections*

Immunofluorescent staining of lung tissue was completed as previously described (21). Lung samples from IPF patients and normal controls were fixed with 10% formalin and embedded in paraffin. For the murine studies, lungs were pressure inflation fixed at 25cm $H_2O$ with 10% buffered formalin for 10 minutes. The trachea was tied off to prevent fluid leakage prior to removal of the lungs from the chest cavity. Mouse lungs were then fixed with 10% buffered formalin in a conical at room temperature overnight, rinsed with $dH_2O$ for 15 minutes, dissected manually, and stored in a tissue cassette in 70% ethanol for subsequent paraffin embedding and tissue sectioning (Neuro-histology laboratory at the University of Pittsburgh). Prior to staining and analysis, tissues sections were deparaffinized and rehydrated by a series of xylene and ethanol washes. Antigen retrieval was completed using sodium citrate buffer, pH 6.0 at 95°C. Tissue was permeabilized with 0.3% Triton X-100 and 1% BSA in PBS and blocked with 2% BSA in PBS. Human and mouse lung sections were stained for: ANT1 (Abcam #ab102032 rabbit polyclonal, 1:500), ANT2 (Abcam #192410 rabbit polyclonal,

1:100), goat anti-mouse or anti-rabbit Alexa 488, 555 and 647 (Molecular Probes). Control sections were stained with non-immune rabbit IgG (#2729P, Cell signaling). All tissue sections were stained with Hoechst at 10 μg/mL for 10 minutes. Sections were mounted with Prolong Gold (Molecular Probes, ThermoFisher) and cured for at least 24 hr at 4°C prior to imaging. Antibodies used for each experiment are designated in the figures. For p21 tissue staining, mouse lung tissue was stained against p21 (Abcam, #ab107099, 1:500) and Hoechst (10 μg/mL). Nuclear p21 fluorescence intensity was quantified using a macro script in Image J Fiji(38). All images were captured on a Nikon A1R confocal microscope with 60x oil objective.

*Single cell RNA sequencing on human lung samples*

Single cell RNA sequencing data was provided by the Königshoff lab (39) and is publicly available at NCBI's Gene Expression Omnibus with accession number GSE190889 and a GitHub repository (https://github.com/KonigshoffLab/GPR87_IPF_2022). Briefly, lung samples from IPF patients and normal controls were enzymatically digested as previously described (39), enriched EpCAM+ lung cells of the distal lung were separated out, and single cell RNA sequencing was performed on the EpCAM+ lung cells. The sequencing results were analyzed using the Cell Ranger pipeline from 10x Genomics (v3.1.0, STAR v2.5.3a) and the Scanpy package (v1.8.0) (40). Reads were aligned to a hg38 human reference genome (GRCH38.97). Barcodes with less than 400 or more than 20,000 detected transcripts were excluded from the single cell RNA library. Cells with a high proportion of mitochondrial-encoded transcripts were excluded. Cells with high background mRNA contamination were detected using the R library package SoupX (41) and also excluded from analysis. Variable genes were selected and ranked, and 3,426 genes identified as occurring in at least three samples were used as input for principal component analysis. Differential gene expression was calculated following leiden cell clustering and batch alignment with BBKNN (42). UMAPs were generated in Seurat (CITE). UMAPs and dotplots showing gene expression were generated via scanpy's pl.umap() and pl.dotplot() function, respectively. To select genes correlated with SLC25A4, cells were over-clustered as small clusters followed by the calculation of average gene expression and correlation coefficients. The correlation gene list was sorted and selected based on spearman's rank correlation coefficient. Correlation between SLC25A4 gene and the selected senescence genes were calculated and displayed via visuz.stat.corr_mat()

function from bioinfokit toolkit (v2.0.8) (Bedre, R. reneshbedre/bioinfokit: Bioinformatics data analysis and visualization toolkit. https://doi.org/10.5281/zenodo.3698145 (2021)).

*Human cell culture*

The human bronchial epithelial cell line (HBEC3-KT) was a gift from John Minna. HBEKT cells were grown in keratinocyte serum free media supplemented according to the Lonza protocol and filtered over a Nalgene Rapid-Flow 0.2 um aPES membrane (Halgene Nunc, Rochester, NY). The human bronchial epithelial cell line (BEAS-2B) and human adenocarcinoma alveolar basal epithelial line (A549) were obtained from ATCC. BEAS-2B cells were maintained in Dulbecco's modified eagle media (DMEM) with Nutrient Mixture F-12 and supplemented with 5% fetal bovine serum and 1% penicillin/streptomycin. A549 cells were maintained in DMEM supplemented with 5% fetal bovine serum and 1% penicillin/streptomycin. All cell lines were cultured on cell culture treated polystyrene plates and were propagated when 70-90% confluent. Cells were propagated by trypsinization with 0.25% trypsin-EDTA (Gibco #25200-056) and neutralized by trypsin neutralizing solution (Gibco #R-002-100). To induce senescence, A549 cells were treated with 10 μg/mL bleomycin (pharmaceutical grade) over 2 days.

*Targeted gene suppression*

BEAS-2B, HBEC3-KT and A549 cells were grown to 25-50% confluency and then transfected with 150 nM siRNA ON-TARGETplus smart pools (Dharmacon: ANT1 (*SLC25A4*, #L-007485-00-0005), ANT2 (*SLC25A5*, #L-007486-02-0005), and non-targeting control pool (#D-001810-10-05) by lipofection with Lipofectamine 3000 (Invitrogen) using the manufacturer's protocol. Briefly, lipid-siRNA complexes were formed in Opti-MEM Reduced Serum Media (ThermoFisher) over 20 minutes at room temperature, and then dripped slowly over the cells. ANT1 was knocked down for 72 hours prior to treatment with bleomycin or any additional experiments. Gene expression levels were verified by RT-qPCR and Western blot (see below).

*Realtime Quantitative PCR*

RNA was isolated from human lung tissue samples, mouse lung tissue samples, and cell culture samples using TRIzol reagent (Invitrogen) according to the manufacturer's protocol. Lung tissues were specifically homogenized in TRIzol using the Bullet Blender Tissue Homogenizer and tubes (NextAdvance Inc.) at a setting of 4 for 5 minutes. Isolated RNA was quantified and assessed for purity using a Nanodrop (Thermo Scientific). RNA was converted to cDNA using High-Capacity cDNA Reverse Transcriptase kit with RNase Inhibitor (Applied Biosystems, 4373966). Real time PCR against genes of interest was performed on cDNA using SYBR green for primer pairs or TaqMan probes conjugated to fluoresceine (FAM) (Thermo Scientific). Each sample using primer pairs was measured at least in duplicate. Real time PCR Taqman probes were Hs00154037_m1 (*SLC25A4*), Hs00854499_g1 (*SLC25A5*), Hs02786624_g1 (GAPDH), Mm01207393_m1 (*Slc25a4*), Mm00846873_g1 (*Slc25a5*), Mm99999915_g1 (*Gapdh*). Each sample using TaqMan probes was measured in triplicate. Relative fold change was calculated by normalizing to GAPDH. Error is calculated as positive or negative error = 2^(Fold change ± √(SEM of ACTB^2 + SEM of gene^2)).

*Western blot*

Human and mouse lung homogenates for were prepared in lysis buffer made from 1x RIPA buffer (Pierce #89900) supplemented with 1x RNAase-free DNAase (ThermoScientific #EN0521) and 1x Halt protease and phosphatase inhibitor cocktail (ThermoScientific #1861281) using the Bullet Blender Tissue Homogenizer (NextAdvance Inc.). For the *in vitro* studies, cells were treated with 0.25% trypsin-EDTA (Gibco #25200-056) for 3-5 minutes, and the trypsin was neutralized with trypsin neutralizing solution (Gibco #R-002-100). Cells were harvested at 10000 g for 10 minutes and the pellets were stored at -20 °C for later. Frozen cell pellets were re-suspended in lysis buffer and lysed by sonication or by multiple rapid freeze thaw cycles in liquid $N_2$ followed by aspiration with a pipette. Gross protein concentrations were determined by Pierce Rapid Gold bicinchoninic acid (BCA) assay (ThermoScientific #A53226). For the Western analysis, at least 10 μg of protein was run on a stain-free 10% SDS-PAGE gel and transferred to a PVDF membrane (Bio-Rad #162-0177). Blots were incubated with primary antibody for at least 16 hours at 4 °C, washed three times for 10 minutes, and then incubated with secondary antibody for 1 hr at room temperature. Primary antibodies used were targeted against p16 INK4a (Thermo MA5-1704), p21 Waf1/Cip1 (Cell Signaling 2947S), and p53 (Cell Signaling 2527S). Secondary antibodies used were Invitrogen GOXRB or GOXMO HRP high XADS (#A16110 and #A16078). Blots were developed using SuperSignal West Pico PLUS Chemiluminescent Substrate (ThermoScientific #34580) and imaged using a ChemiDoc XRS+ (Bio-Rad) detector.

*Senescence beta-galactosidase stain*

HBEC3-KT cells were stained for beta-galactosidase using the Senescence Beta-Galactosidase Staining Kit (Cell Signaling Technology) according to the manufacturer's instructions. Briefly, the cells were washed twice with PBS and then subsequently fixed to the plate with fixative solution. The beta-galactosidase staining solution was made fresh and the pH was adjusted to 6.0 using a freshly calibrated pH meter. Cells were then washed twice with PBS and stained overnight at 37°C in a dry incubator without $CO_2$. The plate was wrapped in paraffin to prevent evaporation during the staining process. The next morning, the staining solution was removed and replaced with PBS. Percent staining was quantified by manual counting.

*Bleomycin and asbestos mouse models*

All animal experiments were performed in accordance with the Institutional Animal Care and Use Committee (IACUC) at the University of Pittsburgh. Animals were housed according to standard housing criteria. Mice strains are under a C57BL/6J background and genotyped to ensure they are wildtype for the Nnt gene, as the C57BL/6J strain is known to carry a mutation in this gene that can affect mitochondrial function. ANT1 null mice were generated from sperm graciously gifted to us from Douglas Wallace at the University of Pennsylvania Children's Hospital. Male and female mice (10-12 weeks of age, n = 6-10 per group) were treated with intratracheal bleomycin sulfate (0.05U in 60ml of PBS for a 25g mouse with dose adjustments per mouse) or saline control, or intratracheal crocidolite asbestos (0.1mg in sterile saline, from the National Institute of Environmental Health & Sciences) or titanium dioxide (inert control particulate, 0.1mg in sterile saline). These doses of bleomycin and asbestos have previously been shown to consistently induce pulmonary fibrosis in mice of a comparable genetic background with a <10% mortality rate. Mice were closely monitored for signed of weight loss or distress following treatment. Partial or incomplete treatments were noted. Weights were obtained daily for 7 days, and then every 2-3 days following until harvest. Mice demonstrating >20% weight loss were sacrificed prior to the planned harvest date. At 28 days post-treatment, mice were euthanized by $CO_2$ inhalation. The lungs were then ligated, excised, and fixed in formalin for 24 hr before embedding in paraffin (as described below). In a subset of animals, lungs did not undergo fixation and instead were excised and placed directly in liquid nitrogen for protein or RNA isolation.

*Bronchoalveolar lavage and leukocyte differentials*

Bronchoalveolar lavage (BAL) was performed twice on sacrificed mice by slowly injecting and recovering 800 μL of PBS into the lungs through a needle catheter inserted into the trachea.

*Lung Tissue Histology*

Following the BALs during the mouse harvest, Histological sections were stained with standard hematoxylin and eosin (H&E) stain using standard protocols. Collagen was detected by trichrome stain with aniline blue.

*Tissue scoring*

Mouse lung tissue slices stained with H&E were manually scored to quantify the severity of fibrosis (fibrosis index) and inflammation (interstitial PMN index and alveolar macrophage index), as previously published (43). The entire lung was scored at 40X magnification excluding large airways and blood vessels when possible.

*Hydroxyproline assay*

Hydroxyproline assay was completed as previously published (43). Whole lung tissue was dried in vacuoles in a 110 °C oven for 48 hr, then dissolved in 6M HCl under nitrogen gas or another 24 hr at 110 °C. Samples were dried for 24 hr, reconstituted with PBS for 1 hour at 60 °C, then analyzed for hydroxyproline as previous described (43). Briefly, samples were processed with chloramine-T solution for 20 minutes and 3.15M perchloric acid for another 5 minutes. Samples were dispensed onto a 96 well plate and P-dimethylamino-benzaldehyde solution was added. After 20 minutes, absorbances at 557 nm for standards of 4-hydroxy-L-proline and samples were read in a 96 well plate-reader spectrophotometer (SpectraMax plus, Molecular Devices).
